# Masitinib limits neuronal damage, as measured by serum neurofilament light chain concentration, in a model of neuroimmune-driven neurodegenerative disease

**DOI:** 10.1101/2024.03.07.583695

**Authors:** Olivier Hermine, Laurent Gros, Truong-An Tran, Lamya Loussaief, Kathleen Flosseau, Alain Moussy, Colin D. Mansfield, Patrick Vermersch

## Abstract

**Background:** Masitinib is an orally administered tyrosine kinase inhibitor that targets activated cells of the innate neuroimmune system. We have studied the neuroprotective action of masitinib on the manifestations of experimental autoimmune encephalitis (EAE) induced axonal and neuronal damage. EAE is a model of neuroimmune-driven chronic neuroinflammation and therefore highly relevant to masitinib’s mechanism of action in neurodegenerative diseases. Importantly, neuronal damage, or prevention thereof, can be rapidly assessed by measuring serum neurofilament light chain (NfL) concentration in EAE-induced mice.

**Methods:** EAE induction was performed in healthy female C57BL/6 mice via active MOG_35-55_ peptide immunization. Treatments were initiated 14 days post EAE induction. On day-0, 39 mice with established EAE symptoms were randomly assigned to 3 treatment groups (n=13): EAE control, masitinib 50 mg/kg/day (M50), and masitinib 100 mg/kg/day (M100). Treatment started on day-1 and ended on day-15. Blood samples were collected at day-1, day-8 (via tail vein sampling) and day-15 (via intracardiac puncture). Assessments included quantification of serum NfL levels along the disease duration, cytokine quantification at day-15, and clinical assessments.

**Results:** Masitinib treatment significantly (p<0.0001) limited NfL production with respect to control; specifically, relative change in serum NfL concentration at day-8 was 43% and 60% lower for the M50 and M100 groups, respectively. Likewise, for the assessment of absolute serum NfL at day-8 and day-15, there was a significantly lower NfL concentration for masitinib treatment as compared with control. Furthermore, EAE mice treated with masitinib showed significantly lower concentrations of several well-established pro-inflammatory cytokines relative to control at day-15. A beneficial effect of masitinib on functional performance was also observed, with both M50 and M100 groups showing significantly less relative deterioration in grip strength at day-15 as compared with control (p<0.001).

**Conclusion:** This study is the first demonstration that masitinib, a drug that targets the innate as opposed to the adaptive neuroimmune system, can lower serum NfL levels, and by extension therefore, neuronal damage, in a neuroimmune-driven neurodegenerative disease model. Overall, findings indicate that masitinib has a neuroprotective effect under conditions of chronic neuroinflammation and therefore plausible disease-modifying activity across a broad range of neurodegenerative diseases.

## BACKGROUND

The measurement of neurofilament light chain (NfL) in biological fluids has been proposed for monitoring the therapeutic effect of drugs aimed at reducing axonal damage. NfL are cytoskeletal proteins that are highly specific for neurons in both the central nervous system (CNS) and the peripheral nervous system. NfL in cerebrospinal fluid (CSF) or the bloodstream is therefore indicative of axonal lesions and/or degeneration and elevated NfL levels are associated with traumatic brain injuries or neurodegenerative diseases (NDD), including amyotrophic lateral sclerosis (ALS), multiple sclerosis (MS), and Alzheimer’s disease (AD). Although this non-specificity limits the use of NfL as a diagnostic biomarker, a growing body of literature shows that because the level of free NfL in serum/plasma directly reflects neuronal damage within the CNS, it can be used as a reliable and easily accessible marker of disease intensity and/or activity across a variety of neurological disorders. As such, serum/plasma NfL is being widely touted as a future biomarker for prognosis, progression, and early detection of general neurodegenerative processes, as well as for monitoring the response of disease-modifying treatment [Gaetani 2019; Palermo 2020].

In MS, clinical relapses and new/active MRI lesions (i.e., CNS injury caused by focal inflammation) results in the release of NfL. Thus, elevated levels of NfL are seen in the CSF and serum of patients with active MS and clinically isolated syndrome. In short-term prognosis studies, baseline serum NfL levels have been associated with worsening of the expanded disability status scale (EDSS) in the first year, number of relapses, and progression of brain atrophy, while in long-term prognosis studies, both baseline and longitudinal measures of NfL levels have been linked to greater cerebral MRI-based brain atrophy [Arroyo Pereiro 2023; Gaetani 2019]. The potential role of NfL as a promising long-term disability prognostic marker has also been demonstrated, with baseline serum NfL levels identified as the most efficient marker for distinguishing aggressive RRMS (defined as patients with an EDSS score of ≥ 6 at ≤ 15 years of follow-up) from benign RRMS (defined as patients with an EDSS score of ≤ 3 at ≥ 10 years of follow-up) [Arroyo Pereiro 2023]. Finally, CSF and serum NfL has been shown to significantly decrease in patients with active forms of MS following 12 months of treatment with disease-modifying therapies that target immune-mediated CNS injury, providing proof-of-concept for NfL as a marker of response to therapy in NDDs [Gaetani 2019].

A growing body of scientific evidence supports the use of NfL as a prognostic biomarker in the broad ALS population, and a risk/susceptibility biomarker among a subset of SOD1 pathogenic variant carriers [Benatar 2024]. Notably, it has been shown that the baseline level of NfL in blood and CSF correlates with the speed and severity of ALS progression, with levels remaining relatively stable over time and disease stage [Verde 2021; Bjornevik 2021; Witzel 2024]. Hence, serum NfL may one day provide a more objective and reliable pharmacodynamic biomarker for therapeutic trials than methods currently used for patient stratification and enrichment; e.g., the ALSFRS-R progression rate (ΔFS).

In patients with AD and mild cognitive impairment (MCI) AD, there is a correlation between plasma NfL concentration and cognitive impairment, MRI hippocampal volume loss and brain atrophy. Moreover, higher plasma NfL levels predict faster cognitive deterioration and a higher rate of brain atrophy and hypometabolism in MCI patients over time [Palermo 2020]. The potential predictive biomarker utility of serum NfL was also demonstrated in familial Alzheimer’s disease, with NfL dynamics predicting disease progression and brain neurodegeneration at the early presymptomatic stage of the disease [Preische 2019].

Evidently therefore, NfL may become an indispensable biomarker for MS, ALS and AD to assess neuronal alterations. Indeed, regulatory proof of potential for this biomarker has now been established through the FDA approval of tofersen for ALS, based on that drug’s ability to lower blood levels of NfL [FDA 2023; Meyer 2023]. Importantly, NfL can serve as a biomarker of a drug’s ability to produce a neuroprotective effect across a broad range of NDD indications, and any meaningful claim of clinical benefit is expected to be supported by a response in NfL.

Masitinib is a tyrosine kinase inhibitor that selectively inhibits c-Kit, platelet-derived growth factor receptor (PDGFR), colony-stimulating factor 1 receptor (CSF1R), LYN, and FYN, in the sub-micromolar range, and which is capable of accumulating in the CNS at therapeutically relevant concentrations [Dubreuil 2009; Trias 2016]. Masitinib also inhibits cellular events mediated by activation of these receptor kinases, with its neuroprotective action resulting predominately from dual targeting of mast cell and microglia/macrophage activity and subsequent remodeling of the neuronal microenvironment. Microglia, macrophages and to a lesser extent mast cells, are types of innate immune cells that are present in the CNS and peripheral nervous system, and which are increasingly recognized as being involved in the pathophysiology of NDDs [Lin 2023; Mado 2023; Kamma 2022; Sandhu 2021; Muzio 2021; Harcha 2021; Pinke 2020; Hagan 2020; Schwabe 2020; Long 2019; Jones 2019; Skaper 2018; Hendriksen 2017; Shaik-Dasthagirisaheb 2016].

To date, masitinib has demonstrated neuroprotective benefits in three challenging NDDs; namely, mild-to-moderate AD, progressive forms of MS, and ALS, both in terms of preclinical models and clinical phase 2b/3 studies [Dubois 2023; Vermersch 2022; Mora 2021; Mora 2020; Kovacs 2021; Trias 2020; Li 2020; Harrison 2020; Trias 2018; Trias 2017; Trias 2016]. However, none of these studies performed fluid-based biomarker analysis. As such, evidence to support modification of underlying disease processes, and in particular neuroprotection resulting in the reduction of neuronal damage, is lacking. As an initial step to address this gap in our knowledge, we have studied the neuroprotective action of masitinib treatment on the manifestations of experimental autoimmune encephalitis (EAE) induced axonal and neuronal damage, using the MOG 35–55 peptide-induced EAE model. EAE is a CNS model of the neuroimmune system (including components of the innate immune system such as mast cells and microglia) that mimics aspects of MS and more generally the features of chronic neuroinflammation, which is a common pathological characteristic of most NDDs [Zhang 2023; Kwon 2020]. In EAE-affected mice, persistent axonal damage occurs from the early disease phase, as revealed by analysis of NfL leakage into the bloodstream along the disease duration, with clinical symptoms typically seen 12 to 16 days after disease induction (depending on EAE induction protocol and age of mice). Hence, axonal and neuronal damage, and prevention thereof, can be rapidly assessed by measuring serum NfL concentration in EAE-induced mice, along the disease duration.

## MATERIALS AND METHODS

### EAE model and negative (non EAE) control animals

A total of 60 female C57BL/6 mice, aged 9-11 weeks old, were purchased from the Charles River Laboratories (Saint-Germain-Nuelles, France). Animals were housed in the pathogen-free animal facility of the Center for Exploration and Experimental Functional Research (CERFE, Evry, France), according to animal protocols approved by the European Directive 2010/63/EU and under valid experimental authorization issued by the French Research Ministry: APAFIS #45222-2023102009547945 v2. All the mice were housed in individually-ventilated cages of standard dimensions (maximum occupancy of 6 animals) in an environmental monitored area (temperature and relative humidity) and with sterilized litter, standard food and water ad libitum, at room temperature under a 14:10 h light-dark cycle. Animals were acclimated to the study conditions for a period of 7 days prior to experimentation. Animals following the EAE protocol received diet supplementation (Dietgel® Recovery capsules) with their food pellets. No special procedure was followed for negative (non EAE) control mice. This experimental procedure was approved by the French experimental animal ethics committee n°051 (CERFE, approval D91228107) under the number 2023-011-B.

### EAE model induction

EAE induction was performed in 47 healthy female mice (aged 10-12 weeks old) via active immunization using an EAE induction kit by Hooke Lab (MOG 35-55/CFA emulsion PTX, ref. EK-2110), administered subcutaneously at two sites (i.e., on midline of upper and lower back) at 0.1 ml/site (i.e., a total of 0.2 ml per mouse). Antigen (MOG 35-55) emulsion was ready-to-use and was kept on ice during the entire induction period. Immediately prior to injection, the pertussis toxin solution (PTX) was prepared by diluting the stock solution in cold PBS (2-8°C) to obtain a final concentration of 110 ng/ul. The solution was gently mixed by inverting (i.e., no vortexing) and kept on ice until used. The PTX solution was administered intraperitoneally at 0.1 mL/dose, 2 hours after antigen emulsion administration. This PTX procedure was repeated 24 hours later. The remaining 13 healthy mice were assigned to a negative (non EAE) control group. The following limit points were defined to euthanize any given animal if they were reached: general comportment alteration, clinical score of strictly greater than 3, or body weight loss of at least 20% at any time compared to baseline weight.

### Experimental groups and blood sampling

Treatments were initiated 14 days post EAE induction. On Day 0 (D0), 39 mice with established EAE symptoms were randomly assigned to 3 treatment groups, comprising 13 mice per group (namely, EAE [vehicle] control, masitinib at 50 mg/kg/d, and masitinib at 100 mg/kg/d). Stratification of these EAE-induced mice was performed based on rotarod performance, this parameter providing and earlier and more objective indication of EAE onset than clinical signs alone, thereby ensuring homogenous clinical functionality across groups at baseline. Additionally, a negative (non EAE-induced) control group consisted of 13 healthy mice, without EAE induction and without any clinical signs. Treatment started on D1 and ended on D15. The test items, masitinib (AB Science, France) or its vehicle (NaCl 0.9%), were dosed twice a day (i.e., at 25 mg/kg or 50 mg/kg for the total daily doses of 50 mg/kg/d or 100 mg/kg/d) for 14 consecutive days by oral gavage applicator with at least 6.5 hours between the 2 daily doses. Blood samples were collected from all animals once a week during the treatment period at D1, D8 and D15. For intermediate blood sampling (D1 and D8 after the morning dose), up to 40 µl of whole blood was collected from each animal via tail vein sampling using a capillary coated with heparin. These intermediate samples were pooled according to treatment group and stored overnight at +4°C prior to serum separation via centrifugation at 5000 rpm for 10 minutes at +4°C. For terminal blood sampling on D15, intracardiac puncture was performed under general anesthesia with a solution of ketamine at 100 mg/kg (Ketamine®) and xylazine at 10 mg/kg (Rompun® 2%). At least 500 µl of whole blood was collected from each animal and stored for at least 2 hours at +4°C prior to serum separation via centrifugation at 1500 rpm for 20 minutes at +4°C. Serum samples were stored at -80°C until analysis.

### Serum biomarker assessments

NfL levels in serum samples from EAE mice and negative controls were quantitated using the R-PLEX Human Neurofilament L assay (Catalog # K1517XR-2, Meso Scale Discovery), which employs advanced Meso Scale Discovery® (MSD) electrochemiluminescence (ECL)-based detection technology. Sera pools samples from D1 and D8 were diluted 2-fold, 4-fold and 8-fold, and were run in duplicates. Sera pools from D15 were diluted 4-fold, 8-fold and 16-fold, and also run in duplicates. Individual sera from D15 were diluted 10-fold. The volume of original sample used was 25 μL of serum per replicate for the NfL quantification and 50 µL of serum per replicate for the cytokine quantification. Because of the larger sample volumes required for cytokine detection, this evaluation was conducted only for D15 intracardiac samples. Cytokines were quantified in serum samples using MSD’S V-PLEX mouse cytokine panel 1 (K15245D) and V-PLEX Proinflammatory Panel 1 Mouse Kit (K15048D-1), according to the manufacture’s instruction, for IFN-gamma, IL-1beta, IL-6, KC/GRO, IL-10, IL-12p70, IL-2, TNF-alpha, IL-33, IL-17A/F, MIP-2. The calibrators supplied with kits were used as standards, calibration curves being constructed using 8 calibrators with concentrations ranging from 0 to 50,000 pg/mL for NfL and from 0 to 27900 pg/mL for cytokines (depending on the cytokine tested). Analysis was done in the MSD Discovery Workbench software, with signal from calibrators fit to standard curves, which were then used for calculation of NfL and cytokine concentration in the samples.

### Clinical assessments

Body weight of each animal was recorded twice per week over the study duration. Each animal was observed twice a day over the study duration with any change in behavior, physical appearance or clinical signs recorded. Health status and disease progression were evaluated daily using an EAE specific clinical scoring method, with scores and clinical observations related as follows: score of 0 – no obvious changes in motor function compared to non-immunized mice; 0.5 – tip of tail is limp; 1.0 – no signs of tail movement observed; 1.5 – limp tail and hind leg inhibition; 2.0 – limp tail and weakness of hind legs or poor balance; 2.5 – limp tail and dragging of hind legs; 3.0 – limp tail and complete paralysis of hind legs; 3.5 – limp tail and complete paralysis of hind legs in addition to no righting reflex (euthanasia recommended); 4.0 – limp tail, complete hind leg, and partial front leg paralysis (euthanasia recommended); 4.5 – complete hind and partial front leg paralysis with no movement around the cage (euthanasia recommended); 5.0 – dead or mouse is spontaneously rolling in the cage (euthanasia recommended).

The effect of masitinib on motor coordination and muscle tone was evaluated by the rotarod test and grip strength test, with animals tested under non-blinded conditions. Disease progression impairs motor coordination/muscle tone, thereby decreasing the time (seconds) an animal can remain on a rotating rod or decreasing the grip force (forelimb and hindlimb), recorded on a strain gauge in Newtons (N). Both tests were performed twice a week, in the morning, at least 1 hour after starting the light circadian cycle. Mice were tested on the rotating rod with a starting speed of 4 rotations per minute (rpm) and acceleration up to the maximum speed of 40 rpm within 300 s (i.e., a single acceleration ramp of 1 rpm every 8.3 s). Each mouse was tested in 3 consecutive trials, with 2–5 minutes between trials, and the best score retained for analysis. Grip strength was obtained by pulling the animal backwards along the platform until the animal’s paws grab the mesh grip piece on the push-pull gauge. The animal was gently pulled backwards with consistent force by the experimenter until it released its grip. Again, each mouse was tested in 3 consecutive trials and the best score retained for analysis.

### Data analysis

Relative change in NfL over time was calculated at D8 relative to D1, using pooled tail vein samples according to treatment group. Because D15 blood samples were collected via intracardiac puncture, intergroup comparison was based on absolute NfL and cytokines concentrations at this single, terminal timepoint. Comparison of absolute NfL measurements at D1 served as a check of possible bias in axonal damage at baseline. Likewise, the non-EAE cohort served as a negative control group to verify that EAE-induced treatment groups were showing the expected axonal damage, as indicated by elevated serum NfL concentrations, and expected concomitant deterioration in clinical performance with disease progression. All group values are presented as mean and standard error of the mean (SEM), with statistical comparisons computed using the unpaired t-test. All statistical analyses were performed with GraphPad Prism10. Statistical significance is indicated by an asterisk * for p≤0.05, ** for p≤0.01, *** for p≤0.001, **** for p≤0.0001.

## RESULTS

### Baseline characteristics and EAE disease course

Table 1 shows that at the time of treatment initiation (D1, 14 days post EAE induction), the EAE treatment groups showed a clear deterioration of functional performance (as measured by rotarod test, grip strength and clinical score) and axonal integrity (as measured by serum NfL concentration), when compared with the negative (non-EAE) control group. These observations are consistent with the description of the EAE model. Clinical baseline characteristics were balanced between the masitinib treatment arms but both of these arms appeared to show slightly more severe disease relative to the EAE control group, as evidenced by the latter having a lower NfL concentration, higher function scores, and lower clinical score.

**Table 1:** Baseline characteristics at time of treatment initiation (D1) according to treatment group

| Experimental group | n | Rotarod (s) | Body weight (g) | Grip strength (N) | Clinical score | Serum NfL (ng/ml) | D1 Relative NfL* |
| --- | --- | --- | --- | --- | --- | --- | --- |
| Negative [non-EAE] control | 13 | 167.1 ± 15.3 | 19.8 ± 0.3 | 229.4 ± 8.4 | 0 | 3.2 ± 0.04 | N/A |
| EAE [vehicle] control | 13 | 109.4 ± 17.2 | 19.5 ± 0.6 | 197.9 ± 16.1 | 0.35 ± 0.13 | 7.3 ± 0.2 | 1.0 |
| Masitinib 50 mg/kg/d | 13 | 100.4 ± 13.5 | 19.2 ± 0.5 | 172.3 ± 14.8 | 0.58 ± 0.12 | 9.4 ± 0.2 | 1.3 |
| Masitinib 100 mg/kg/d | 13 | 100.2 ± 12.4 | 19.4 ± 0.3 | 183.1 ± 13.0 | 0.58 ± 0.10 | 13.5 ± 0.2 | 1.8 |

Values are expressed as mean ± SEM. \* D1 Relative NfL is the ratio of concentrations with respect to the EAE [vehicle] control group. N/A = not applicable.

One mouse from the EAE control group was euthanized at D7, according to the ethical criteria. All other animals were alive and with a clinical score of less than 3.5 at the terminal timepoint (D15).

### Masitinib slows deterioration of grip strength in the EAE mouse model

The non-EAE cohort served as a negative control group to verify that EAE-induced animals, in particular the EAE control group, were experiencing a deterioration in clinical performance consistent with the description of the EAE model. All clinical assessments, including average relative changes in grip strength, clinical score, rotarod performance, and body weight, showed a significant (p <0.001) worsening during the treatment period (D1 to D15) for the EAE control group with respect to the non-EAE cohort (which conversely remained stable), confirming development of an EAE phenotype.

A beneficial effect of masitinib on functional performance was observed in terms of grip strength. While there was a significant deterioration in relative grip strength over the 15-day treatment period for the EAE control group as compared with the negative control group, the grip strength capabilities of masitinib treated mice initially deteriorated but then recovered to their pretreatment (D1) level by D15. Indeed, both the 50 and 100 mg/kg/d masitinib groups showed significantly less relative deterioration in grip strength at D15 with respect to the EAE control group (p <0.001) (Figure 1).

**Figure 1:**
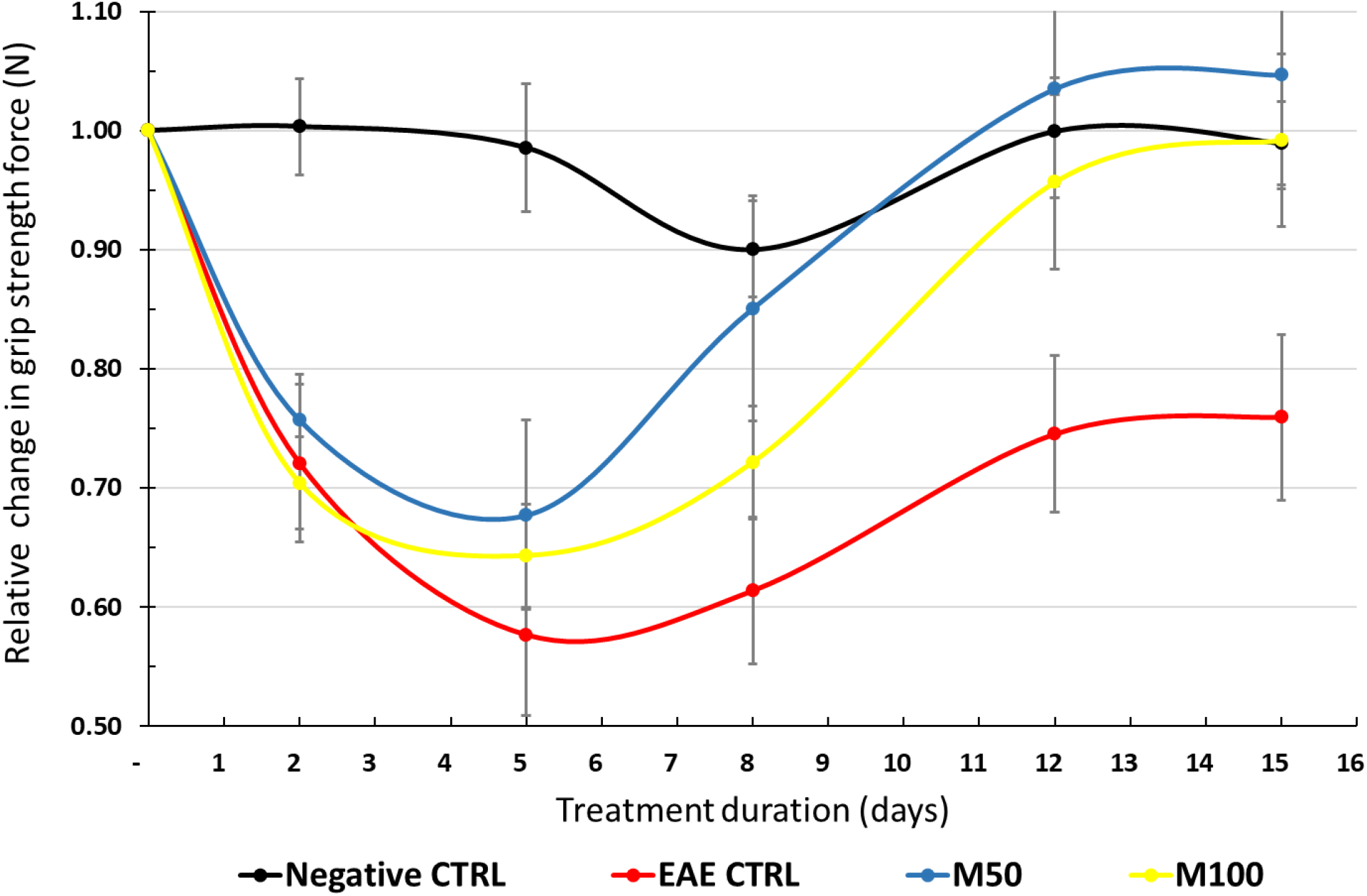
Average relative deterioration in grip strength force (N) over treatment period, according to treatment group CTRL = EAE control group. M50 = Masitinib 50 mg/kg/d. M100 = Masitinib 100 mg/kg/d. N = Newtons. Relative grip strength normalized to value at start of treatment period (D0). Values are expressed as mean ± SEM.

Conversely, there was no discernable difference between EAE control and masitinib treated groups for any of the other clinical assessments over the treatment period.

### Masitinib reduces serum NfL concentration in the EAE mouse model

As expected, all EAE groups showed an increased concentration of serum NfL with disease evolution, while that of the negative control group remained stable or decreased slightly. However, assessment of relative change in serum NfL concentration over a duration of 8 days (pooled tail vein sampling) showed that masitinib treatment significantly reduced NfL production in EAE mice with respect to the EAE control group (Table 1, Figure 2). The relative increase at D8 with respect to D1 was 3.5-fold for the EAE control group, which was a significantly greater relative change than either the 50 mg/kg/d masitinib group (2-fold increase at D8 with respect to D1, p <0.0001), or the 100 mg/kg/d masitinib group (1.4-fold increase at D8 with respect to D1, p <0.0001). This corresponds to a relative reduction in serum NfL concentration of 43% and 60% for the masitinib 50 mg/kg/d and masitinib 100 mg/kg/d groups relative to the EAE control group, respectively. Pairwise comparison of the masitinib groups also showed a dose-dependent effect with a significantly smaller relative change for masitinib 100 mg/kg/d as compared with masitinib 50 mg/kg/d (p <0.0001).

**Figure 2:**
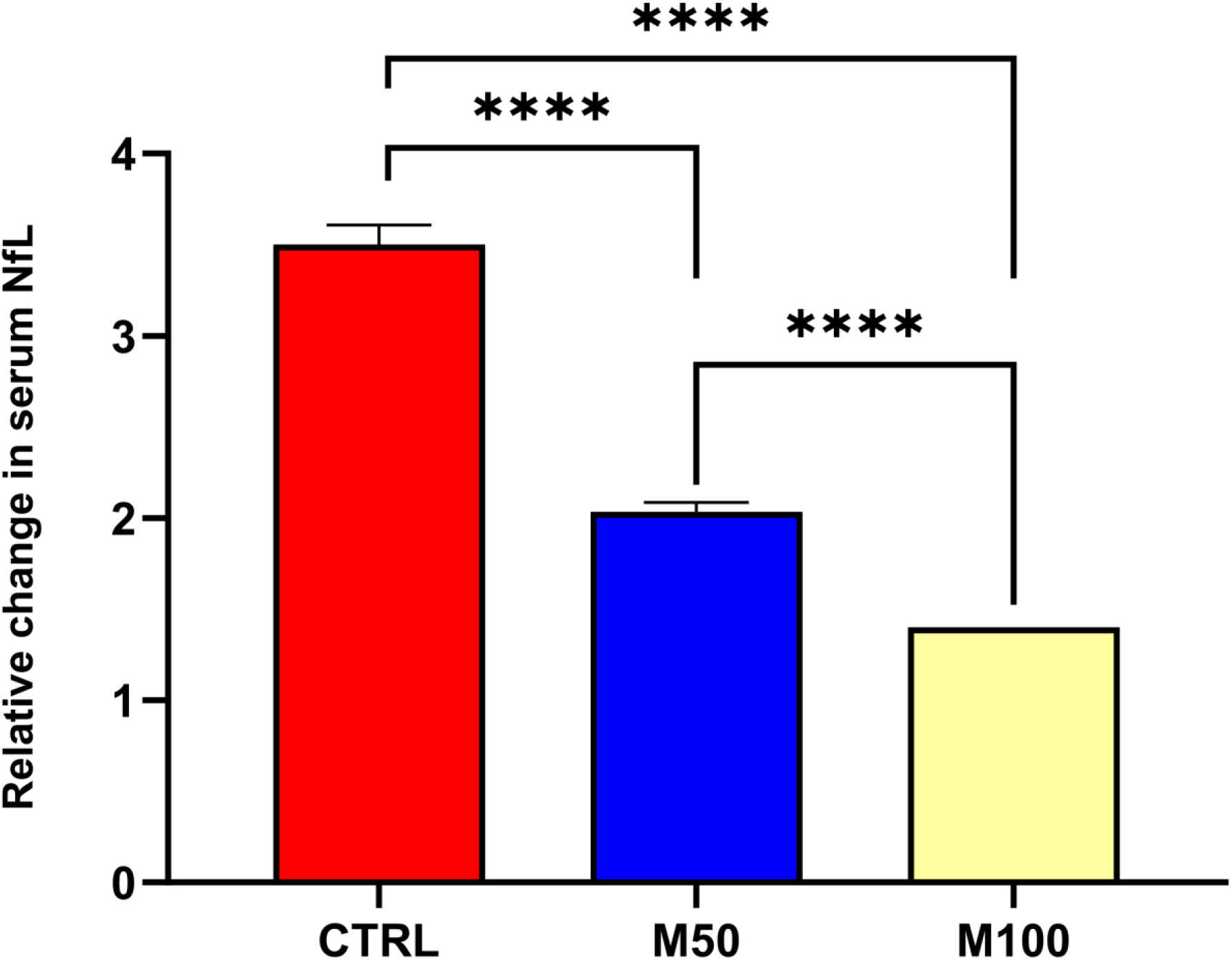
Relative change from baseline at Day-8 in serum NfL concentration (pooled tail vein sampling), according to treatment group CTRL = EAE control group. M50 = Masitinib 50 mg/kg/d. M100 = Masitinib 100 mg/kg/d. Values are expressed as mean ± SEM. Statistical significance (unpaired t-test) is indicated by an asterisk * p<0.05, **** for p<0.0001

Consistent with what was seen for the relative change in serum NfL over 8 days, the absolute concentration at D8 was significantly lower by approximately 25% for both masitinib treatment groups (50 and 100 mg/kg/d) relative to the EAE control group (p <0.0001) (Table 2, Figure 3). By comparison, at baseline (D1) the EAE control group NfL concentration had started at a lower absolute concentration than both masitinib groups (Table 1).

**Figure 3:**
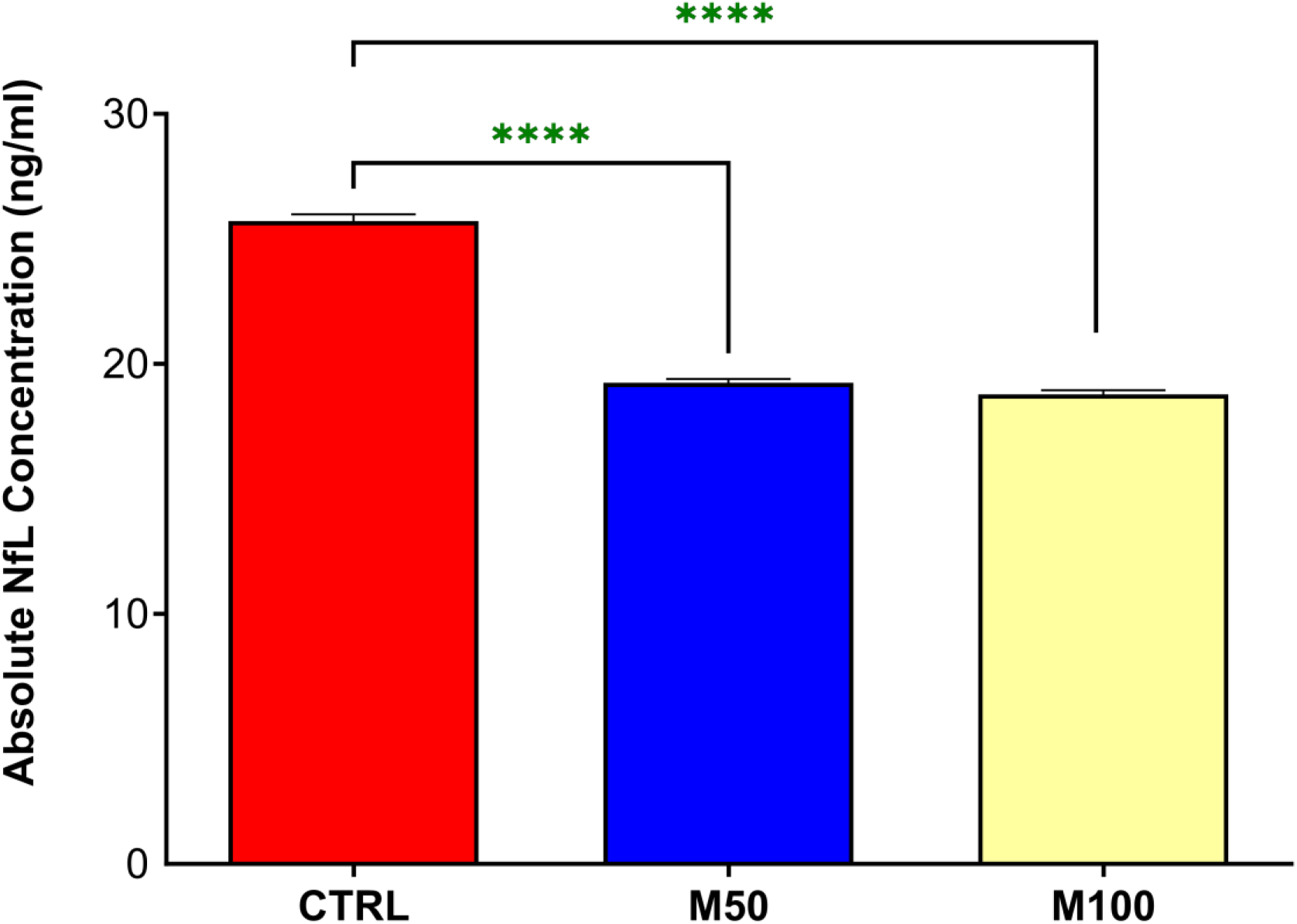
Day-8 absolute serum NfL concentrations (pooled tail vein sampling), according to treatment group CTRL = EAE control group. M50 = Masitinib 50 mg/kg/d. M100 = Masitinib 100 mg/kg/d. Values are expressed as mean ± SEM. Statistical significance (unpaired t-test) is indicated by an asterisk * p<0.05, **** for p<0.0001

**Table 2:** Assessment of serum NfL concentration (pooled tail vein sampling) – Relative change in serum NfL concentration over time (D8 relative to D1), and D8 absolute concentration

| Experimental group | n | Relative Change<br>(D8:D1) | p-value<sup>†</sup> | Absolute NfL at D8<br>(ng/ml) | D8 Relative<br>Difference NfL * |
| --- | --- | --- | --- | --- | --- |
| EAE [vehicle] control | 13 | 3.5 | N/A | 25.7 ± 0.28 | N/A |
| Masitinib 50 mg/kg/d | 13 | 2.0 | < 0.0001 | 19.2 ± 0.16 | -25% |
| Masitinib 100 mg/kg/d | 13 | 1.4 | < 0.0001 | 18.8 ± 0.18 | -27% |

Values are expressed as mean ± SEM. P-value with respect to the EAE [vehicle] control group. † unpaired t-test with respect to the EAE [vehicle] control group. \* D8 Relative Difference NfL is the proportional difference in concentrations with respect to the EAE [vehicle] control group. N/A = not applicable.

Likewise, for the assessment of serum NfL at D15 (pooled intracardiac puncture sampling), there was a significantly lower concentration for both masitinib treatment groups (50 and 100 mg/kg/d) as compared with the EAE control group (Figure 4). The treatment effect was more pronounced for the masitinib 100 mg/kg/d group as compared with the masitinib 50 mg/kg/d group, with an average lower concentration than the EAE control group of 26% (p <0.0001) and 6% (p <0.001), respectively, and a significant dose-dependency (p <0.0001) between masitinib groups. Due to the larger quantity of serum collected from intracardiac puncture sampling, it was possible to also assess samples from each individual animal. As seen in Figure 5, there is a large inter-subject spread of NfL concentrations, however, the average values generally reflect the pooled D15 results, with a significantly lower concentration for the masitinib 100 mg/kg/d treatment group relative to the EAE control group (p <0.0037), and a significant dose-dependency (p <0.0086) between masitinib groups.

**Figure 4:**
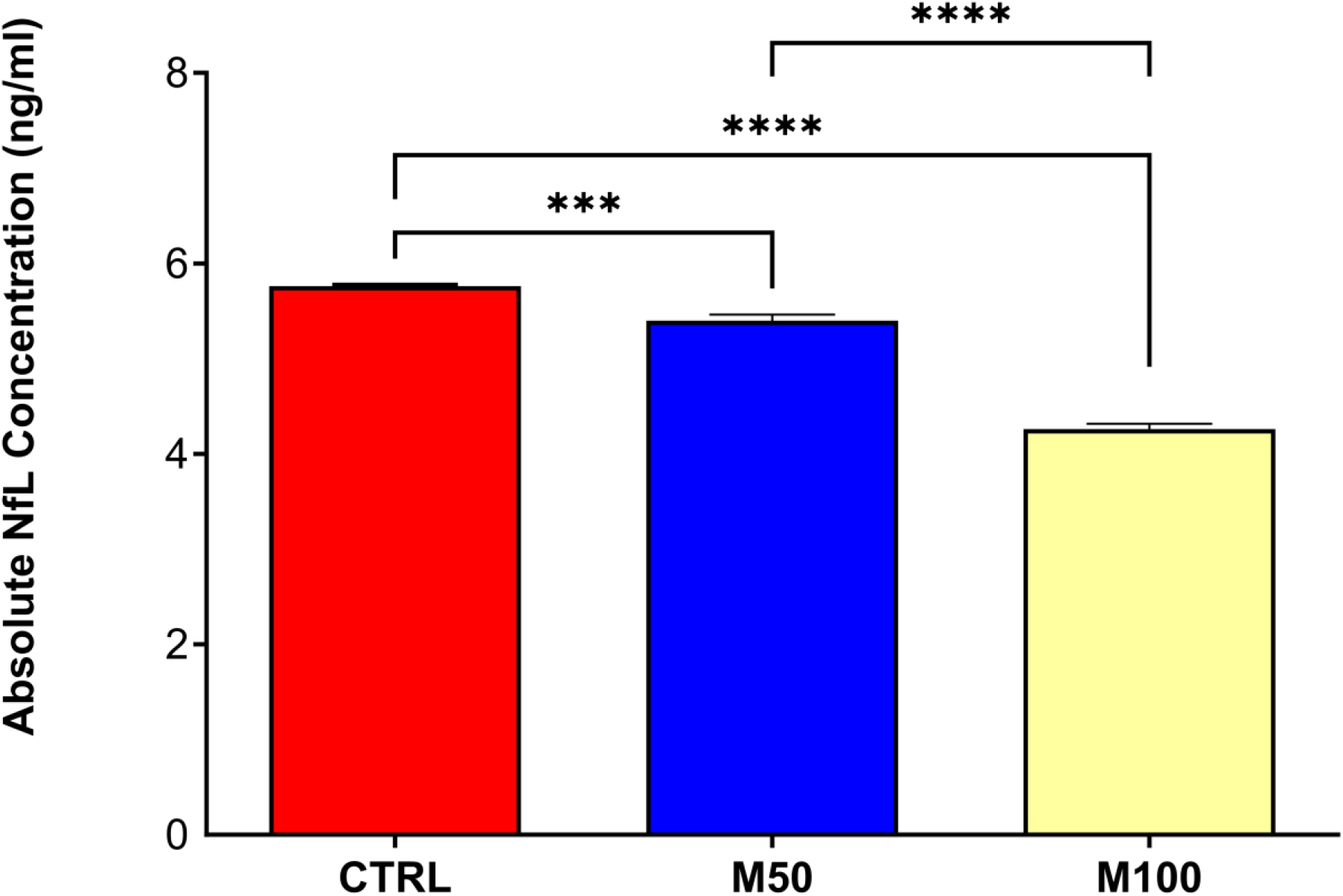
Day-15 absolute serum NfL concentrations (pooled tail vein sampling), according to treatment group CTRL = EAE control group. M50 = Masitinib 50 mg/kg/d. M100 = Masitinib 100 mg/kg/d. Values are expressed as mean ± SEM. Statistical significance (unpaired t-test) is indicated by an asterisk * p<0.05, *** for p<0.001, **** for p<0.0001

**Figure 5:**
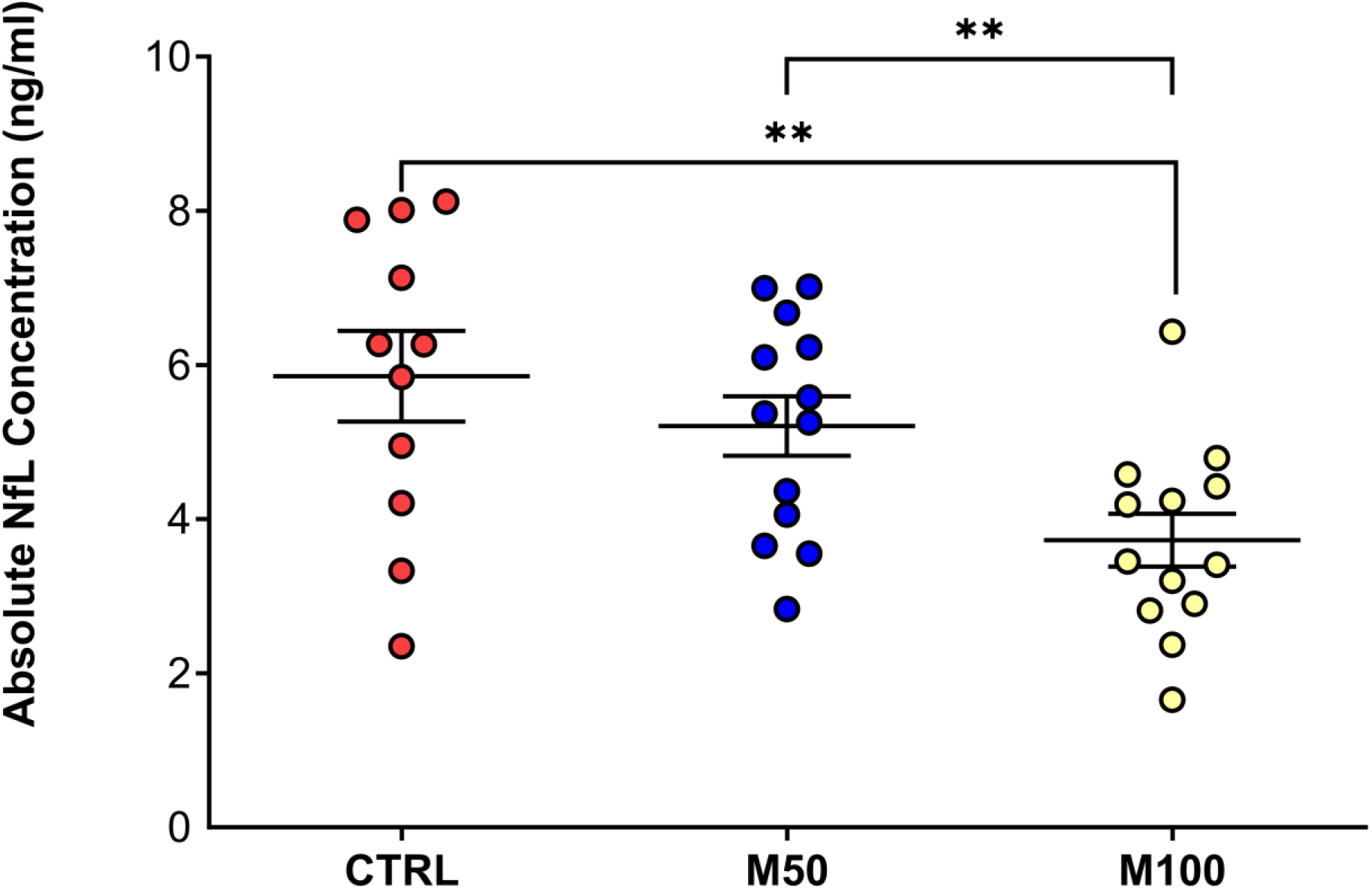
Day-15 absolute serum NfL concentrations from individual animals (intracardiac puncture sampling), according to treatment group CTRL = EAE control group. M50 = Masitinib 50 mg/kg/d. M100 = Masitinib 100 mg/kg/d. Values are expressed as mean ± SEM. Statistical significance (unpaired t-test) is indicated by an asterisk * p<0.05, ** for p<0.01

### Masitinib decreases pro-inflammatory cytokine biomarker concentrations in the EAE mouse model

Cytokines that are important in inflammatory responses and immune system regulation, including those associated with Th1/Th2 pathway biomarkers, were quantified to determine whether masitinib treatment was associated with lower levels of these inflammatory biomarkers. Serum samples from D15, collected using the intracardiac puncture procedure, were used for this analysis because of the larger volume requirements necessary.

Overall, EAE mice treated with masitinib showed significantly lower concentrations of several well-established pro-inflammatory cytokines, i.e., interferon gamma, tumor necrosis factor alpha, human interleukin-1 beta, interleukin-33, KC/GRO, MIP-2, as compared with the EAE control group (Table 3). This is consistent with there being reduced neuroinflammatory activity.

**Table 3:** Relative difference in serum cytokine concentrations (pooled intracardiac puncture sampling) with respect to the EAE control group following 15 days of treatment

| | n | IFNγ | | TNFα | | IL-1B | | KC/GRO | | IL-33 | | MIP-2 | |
| --- | --- | --- | --- | --- | --- | --- | --- | --- | --- | --- | --- | --- | --- |
|  |  | Rel Diff (%) | P | Rel Diff (%) | P | Rel Diff (%) | P | Rel Diff (%) | P | Rel Diff (%) | P | Rel Diff (%) | P |
| **Ctrl** | 11 | N/A |  | N/A |  | N/A |  | N/A |  | N/A |  | N/A |  |
| **M50** | 13 | -40±18 | <0.0001 | -22±5 | <0.0001 | -33±15 | <0.0092 | +2±3 | ns | -82±4 | <0.0001 | -4±3 | ns |
| **M100** | 13 | -34±1 | <0.0001 | -10±1 | <0.0036 | -37±5 | <0.0011 | -15±4 | <0.0004 | -48±14 | <0.0001 | -21±10 | <0.0004 |

M50 = Masitinib 50 mg/kg/d. M100 = Masitinib 100 mg/kg/d. Ctrl = EAE control group. Values are expressed as mean ± SEM. ns = statistically non-significant. P = p-value calculated via unpaired t-test. N/A = not applicable. IFN $\gamma$ = Interferon gamma. TNF $\alpha$ = Tumor necrosis factor alpha. IL-1B = Human interleukin-1 beta. KC/GRO = Keratinocyte chemoattractant/human growth-regulated oncogene. IL-33 = Interleukin-33. MIP-2 = Macrophage inflammatory protein-2.

Interferon gamma (IFNg), also known as immune interferon, is a pro-inflammatory cytokine produced by lymphocytes and is a potent activator of macrophages. It plays physiologically important roles in promoting innate and adaptive immune responses, and is involved in the regulation of anti-inflammatory responses. The absolute concentration of IFNg at D15 was significantly lower for the masitinib 50 mg/kg/d and masitinib 100 mg/kg/d groups relative to the EAE control group by 40% and 34%, respectively (i.e., 1.8 ± 0.6 and 2.1 ± 0.1 versus 3.1 ± 0.2 pg/ml, respectively).

Tumor necrosis factor alpha (TNFa) is an inflammatory cytokine that is mainly secreted by macrophages. The absolute concentration of TNFa at D15 was significantly lower for the masitinib 50 mg/kg/d and masitinib 100 mg/kg/d groups relative to the EAE control group by 22% and 10%, respectively (i.e., 13.4 ± 0.9 and 15.4 ± 0.2 versus 17.1 ± 0.1 pg/ml, respectively).

Human interleukin-1 beta (IL-1b) is a pro-inflammatory cytokine that is produced by macrophages/monocytes during acute inflammation and which contributes to a variety of NDDs. The absolute concentration of IL-1b at D15 was significantly lower for the masitinib 50 mg/kg/d and masitinib 100 mg/kg/d groups relative to the EAE control group by 33% and 37%, respectively (i.e., 0.7 ± 0.2 and 0.6 ± 0.1 versus 1.0 ± 0.1 pg/ml, respectively).

Interleukin-33 (IL-33) is a member of the IL-1 cytokine family and is considered to be crucial for induction of Th2-type cytokine-associated immune responses. Its expression is upregulated following pro-inflammatory stimulation. The absolute concentration of IL-33 at D15 was significantly lower for the masitinib 50 mg/kg/d and masitinib 100 mg/kg/d groups relative to the EAE control group by 82% and 48%, respectively (i.e., 0.8 ± 0.2 and 2.4 ± 0.8 versus 4.7 ± 0.2 pg/ml, respectively).

Keratinocyte chemoattractant (KC)/human growth-regulated oncogene (GRO) (KC/GRO) is a CXC chemokine, which is involved in neutrophil activation and recruitment during inflammation. The absolute concentration of KC/GRO at D15 was significantly lower for the masitinib 100 mg/kg/d group relative to the EAE control group by 15% (i.e., 50.7 ± 3.8 versus 59.6 ± 1.5 pg/ml, respectively).

Macrophage inflammatory protein (MIP-2) is a CXC chemokine, also known as chemokine CXC ligand (CXCL2). MIP-2 affects neutrophil recruitment and activation. The absolute concentration of MIP-2 at D15 was significantly lower for the masitinib 100 mg/kg/d group relative to the EAE control group by 21% (i.e., 9.5 ± 1.6 versus 12.0 ± 0.6 pg/ml, respectively).

Other tested cytokines, i.e., IL-6, IL-17, IL-2, IL-10 and IL-12p70, showed no discernable difference between EAE control and masitinib treated groups.

## DISCUSSION

This study is the first demonstration that masitinib can lower serum NfL levels in a neuroimmune-driven neurodegenerative disease model, with concomitant reduction in pro-inflammatory cytokines and slowing of clinical (EAE) symptoms. Strengths of this study are that it used the well-established MOG 35–55 peptide-induced EAE model of neuroimmune-driven chronic neuroinflammation, which is highly relevant to masitinib’s mechanism of action (i.e., targeting of the innate neuroimmune system and in particular mast cells and microglia). Because chronic neuroinflammation is a common pathological characteristic of most NDDs, these results should have broad applicability across a range of indications. Moreover, data was derived after disease onset (i.e., in a therapeutic setting as opposed to an asymptomatic preventative setting), which is of greater relevance because such models more closely simulate the clinical condition of NDD patients and therefore better represent their therapeutic needs. Limitations of the study included insufficient sample volume to perform both NfL and cytokine assays at multiple timepoints across the study duration, inconsistency in blood collection techniques at different timepoints (tail vein versus intracardiac, which limited availability of comparable data for calculation of relative change from baseline), and a short duration of treatment that may not have allowed for measurable cytokine changes (i.e., due to therapeutic lag). As such, it would be useful to confirm these findings using standardized sample handling techniques, at a schedule that permits longitudinal analysis of both NfL and cytokines, over a longer treatment period. Ideally, findings would also be supported by pathological data showing a correlation between neuronal preservation and NfL concentration, as well as assessment of mast cell activation markers to better discriminate the respective contributions of microglia and mast cells.

Remarkably, masitinib has so far demonstrated clinical benefit in three challenging neurodegenerative disorders. In progressive forms of MS, masitinib administered at 4.5 mg/kg/d showed a statistically significant reduction in cumulative change on EDSS score with masitinib 4.5 mg/kg/d (p=0.0256). This treatment-effect was consistent for both primary progressive MS and non-active secondary progressive patient subgroups. Masitinib also significantly reduced the risk of reaching an EDSS score of 7.0, corresponding to disability severe enough that the patient is restricted to a wheelchair (p=0.0093) [Vermersch 2022]. The rationale to use masitinib in progressive forms of MS is supported by evidence from the scientific literature, showing that progressive MS is driven by activity of the innate immune system, compartmentalized within the CNS [Kamma 2022; Mahmood 2022; Pinke 2020; Stys 2019; Jones 2019; Brown 2018; Luo 2017].

In mild-to-moderate AD, masitinib administered at 4.5 mg/kg/d as an add-on therapy to standard of care significantly slowed cognitive deterioration relative to placebo with a manageable safety profile. More specifically, the between-group difference in the Alzheimer’s disease assessment scale-cognitive subscale (ADAS-cog) change from baseline at week 24 was -2.15 (97.5%CI [-3.48, -0.81]), p=0.0003 [Dubois 2023]. This change is considered clinically meaningful, especially when considering its administration on a background of cholinesterase inhibitors and memantine, a 2-point change being consistent with published recommendations [Vellas 2008] and benchmark ADAS-Cog benefit according to well-established therapies [Birks 2018; Birks 2015; Birks 2006]. The rationale to use masitinib in AD is supported by preclinical evidence demonstrating that the pharmacological action of masitinib in mast cells can restore normal spatial learning performance in a mouse model of AD and promotes recovery of synaptic markers [Li 2020; Lin 2023]. Overall, these findings are consistent with a growing body of evidence implicating mast cells and microglia with the pathophysiology of AD [Leng 2021; Kang 2020; Schwabe 2020; Kwon 2020; Long 2019; Nordengen 2019; Fani Maleki 2019; Jones 2019; Kempuraj 2019; Skaper 2018; Hendriksen 2017; Hansen 2017; Shaik-Dasthagirisaheb 2016].

In ALS, masitinib administered at 4.5 mg/kg/d as an add-on to standard riluzole, significantly slowed functional decline at week 48 relative to riluzole alone by 27% [Mora 2020]. For the prespecified primary efficacy population, there was a between-group difference in ALSFRS-R change from baseline of 3.4 points (95%CI [0.65;6.13]; p=0.016). Moreover, long-term follow-up analysis showed a significantly prolonged survival of 25 months (p=0.0478) and 44% reduced risk of death (hazard ratio 0.56 (95%CI [0.32;0.96]), Cox p=0.036), provided that treatment was initiated at an early stage of disease [Mora 2021]. Masitinib’s mechanism of action in ALS has been well-demonstrated in the preclinical setting, with data showing that masitinib co-targets independent pathological mechanisms in different cell types of the brain, spinal cord and peripheral nervous system components that taken together conserves neuro-muscular function [Kovacs 2021; Trias 2020; Harrison 2020; Trias 2018; Trias 2017; Trias 2016]. Again, these findings are consistent with accumulating evidence indicating that immune dysfunction and neuroinflammation are important pathological characteristics of ALS [Vahsen 2020; Clarke 2020; Jones 2019; Skaper 2018; Iyer 2018; Crisafulli 2018].

Hence, the preclinical biomarker data reported here, taken together with the positive clinical outcomes and growing body of literature described above, support a proposition that modulation of the neuroimmune system via inhibition of microglia and/or mast cell activity, is a valid therapeutic strategy across a broad range of NDD indications. Individually, each of these neuroimmune cells represents a viable target for therapeutic intervention, with the dual-targeting of masitinib making it a particularly attractive treatment option.

## CONCLUSIONS

These results provide further evidence regarding the anti neuro-inflammatory properties of masitinib. Importantly, the observed NfL treatment response supports the assertion that masitinib has a neuroprotective effect under conditions of chronic neuroinflammation and therefore plausible disease-modifying activity in each of the NDDs for which it has shown clinical benefit, i.e. progressive forms of MS, ALS, and mild-to-moderate AD.

## LIST OF ABBREVIATIONS

AD: Alzheimer’s disease
ADAS-cog: Alzheimer’s disease assessment scale-cognitive subscale
ALS: Amyotrophic lateral sclerosis
ALSFRS-R: Revised amyotrophic lateral sclerosis functional rating scale
CI: Confidence intervals
Ctrl: EAE control group
CNS: Central nervous system
CSF: Cerebrospinal fluid
EAE: Experimental autoimmune encephalitis
EDSS: expanded disability status scale
IFNg: Interferon gamma
IL-1B: Human interleukin-1 beta
IL-33: Interleukin-33
KC/GRO: Keratinocyte chemoattractant/human growth-regulated oncogene
M50: Masitinib 50 mg/kg/day treatment group
M100: Masitinib 100 mg/kg/day treatment group
MIP-2: Macrophage inflammatory protein-2.
MMSE: Mini–mental state examination
MOG: Myelin oligodendrocyte glycoprotein
MRI: Magnet resonance imaging
MS: Multiple sclerosis
N/A: Not applicable.
NDD: Neurodegenerative disease
NfL: Neurofilament light chain
SD: Standard deviation
SEM: Standard error of the mean
TNFa: Tumor necrosis factor alpha

## DECLARATIONS

### Ethics approval

All experiments are performed in accordance with the European Directive 2010/63/EU and under valid experimental authorization issued by the French Research Ministry: APAFIS #45222-2023102009547945 v2 (valid for 5 years from October 29th, 2023). This experimental procedure was approved by the French experimental animal ethics committee n°051 (CERFE, approval D91228107) under the number 2023-011-B.

### Authors’ contributions

TAT, LL, KF, and LG contributed to data collection.

OH, PV, AM, LG, LL, KF, and TAT were involved in study conception and design. LL, TAT, LG, and CDM did the data interpretation.

CDM wrote the manuscript with contributions from TAT, LL and LG.

All authors contributed to manuscript draft revisions, provided critical comment, and approved submission for publication.

